# Phylogenetic Analysis of a Bacterial Strain of the Genus *Rothia* Detected in Suspension Culture Cells of *Arabidopsis thaliana* (L.) Heynh as a Member of the Family *Micrococcaceae*

**DOI:** 10.1101/2023.07.27.550883

**Authors:** Sergei Shchyogolev, Lev Dykman, Alexander Sokolov, Oleg Sokolov, Larisa Matora

**Affiliations:** Institute of Biochemistry and Physiology of Plants and Microorganisms, Saratov Scientific Centre of the Russian Academy of Sciences (IBPPM RAS), Saratov 410049, Russia

**Keywords:** *Arabidopsis* suspension culture, bacterial isolate, *Micrococcaceae*, *Rothia*, whole genome test, phylogenetic analysis

## Abstract

We report phylogenetic studies of a bacterial isolate (Isolate SG) recovered from a suspension culture of *Arabidopsis thaliana* (L.) Heynh. In doing this, we use the known results acquired by whole genome sequencing of the DNA of *Micrococcaceae* strains closely related to Isolate SG in the 16S rRNA gene test and we evaluate the intra- and intergeneric taxonomic relationships between them using a set of five whole genome tests (ANI, AAI, MiGA, GTDB-Tk, and AAI-profiler). Quantitative analysis of the clustering of the proteomes of these strains by the average amino acid identity (AAI)-based test showed the need to clarify (with possible renaming) the generic assignment of the strains both within and between the identified monophyletic groups. The need for such reclassification was also shown by the AAI-profiler test (Medlar *et al*., 2018) against the UniProt database (250 million records) with the proteome of *Rothia* sp. ND6WE1A – a strain most evolutionarily similar to Isolate SG. The contradictions in the historically given names of strains and metagenomic objects at the genus and family levels, which were identified by using sets of the genomes and proteomes of the strains related to Isolate SG, can be eliminated with appropriate reclassification of the objects by using quantitative criteria in the AAI-based tests.

## 1. Introduction^1^

In a suspension culture of *Arabidopsis thaliana* (L.) Heynh (All-Russian Collection of Higher Plant Cell Cultures, Timiryazev Institute of Plant Physiology of the Russian Academy of Sciences, Moscow, Russia), we have previously found the presence of a bacterial microflora that did not inhibit plant growth (Sokolov *et al*., 2921). The bacteria were gram-positive, non-acid-resistant, and shaped closely to spheres (diameter, about 1 µm). The analysis results for the isolate’s 16S rRNA gene sequence (GenBank OQ702765.1) (DNA sequencing was done by the Research and Production Company SYNTOL, http://www.syntol.ru) indicate that in the 16S rRNA-based test, the isolate (hereafter Isolate SG) is the most closely related to members of the genus *Rothia*.

*Rothia* is a genus of gram-positive, aerobic, coccoid or bacillary, nonmotile, non-spore-forming bacteria of the family *Micrococcaceae*, phylum *Actinobacteria* (Austin, 2015). To date, 15 *Rothia* species have been identified. *Rothia* are part of the normal microflora of the gastrointestinal tract (especially the oral cavity and stomach) of humans and animals, but they can cause gastric atrophy and intestinal metaplasia and induce opportunistic infections of the upper respiratory tract in immunocompromised people (Nardone and Compare, 2015; Odeberg *et al*., 2023). Various *Rothia* have been recovered from soil, water sources, benthos, rocks, atmosphere, and other sources (Austin, 2015).

Endophytic *Rothia* strains of different species have been isolated from the rhizosphere and tissues of the following plants: *Sphagnum magellanicum* (Opelt *et al*., 2007), *Dysophylla stellata* (Xiong *et al*., 2013), *Hedysarum perrauderianum*, *H. naudinianum* (Torche *et al*., 2014), *Musa acuminate* (Sekhar and Thomas, 2015), *Oryza sativa* (Evangelista *et al*., 2017), *Seidlitzia rosmarinus* (Shurigin *et al*., 2020), *Camellia sinensis* (Borah and Thakur, 2020), *Alnus glutinosa* (Davis *et al*., 2020), *A. incana* (Mercurio *et al*., 2022), *Zea mays* (Pisarska and Pietr, 2015; Elbahnasawy *et al*., 2021), *Vaccinium myrtillus* (Mažeikienė *et al*., 2021), *Santalum album* (Tuikhar *et al*., 2022), *Miscanthus floridulus* (Xiao *et al*., 2023), and *Arabidopsis thaliana* (Sokolov *et al*., 2021). *Rothia* endophytes are inhibitory to several pathogenic fungi, bacteria, parasitic nematodes, and insect larvae, and they can be used as biofertilizers (Bano and Muqarab, 2017; da Silva *et al*., 2018; Asadu *et al*., 2020; Nuaima, 2022). In addition, some *Rothia* have the properties of plant growth promoting rhizobacteria, because they can fix nitrogen (Gtari *et al*., 2012) and produce indole-3-acetic acid, improving plant growth and development (Asyakina *et al*., 2023).

Many *Rothia* genomes have biosynthetic gene clusters that supposedly produce antibiotic nonribosomal peptides, siderophores, and other secondary metabolites that modulate microbe– microbe (quorum sensing) and, potentially, microbe–host interactions (de Oliveira *et al*., 2022). Certain strains produce natural substances with antimicrobial activity, which are planned to be used to treat human and animal infections (Fatahi-Bafghi, 2021). In particular, peptidoglycan hydrolases produced by *Rothia dentocariosa* can be used as highly specific therapeutic agents against nasal pathogens (Stubbendieck *et al*., 2023). Finally, a number of *Rothia* soil bacteria can degrade xenobiotics, specifically phthalate esters and aromatic hydrocarbons (Yastrebova and Plotnikova, 2020; Jia *et al*., 2022).

Analysis of the systematic position of Isolate SG (Sokolov *et al*., 2021) and the phylogenetic studies of *Rothia* members in the 16S rRNA-based test (Xiong *et al*., 2013; Elbahnasawy *et al*.,2021; Tuikhar *et al*., 2022; de Oliveira *et al*., 2022; Fatahi-Bafghi, 2021; Stubbendieck *et al*., 2023; Yastrebova and Plotnikova, 2020; Fan *et al*., 2002; Ko *et al*., 2009) have shown their taxonomic proximity to other *Micrococcaceae*, including *Kocuria*, *Arthrobacter*, *Micrococcus*, and other genera.

Here we report an extended phylogenetic analysis of Isolate SG (Sokolov *et al*., 2021) that uses the known results of whole-genome sequencing of the DNA of the *Micrococcaceae* strains closely related to Isolate SG, and we evaluate the intra- and intergeneric taxonomic relationships between them using a set of quantitative whole genome tests.

## 2. Materials and Methods

As the initial sets of strains, 16S rRNA gene sequences, and genomes, we used the results generated by the 16-based ID tool (Search EzBioCloud Database, https://www.ezbiocloud.net) (Table 1) with the 16S rRNA gene sequence of Isolate SG (GenBank OQ702765). This strain was deposited by us in the Collection of Rhizosphere Microorganisms (http://collection.ibppm.ru), Institute of Biochemistry and Physiology of Plants and Microorganisms, Russian Academy of Sciences (IBPPM RAS) under number IBPPM 684.

**Table 1.** Characteristics of the type strains related to Isolate SG (16S rRNA-based test) (Search EzBioCloud Database, https://www.ezbiocloud.net).

| No. | Species# | Strain | GenBank access code to the record for 16S rRNA | GenBank access code to the record for the genome assembly | Pair-wise identity I (%) |
| --- | --- | --- | --- | --- | --- |
| 1 | *R. amarae* * | JCM 11375<sup>T</sup> | AY043359 | GCA_001683935.1 | **99.8** |
| 2 | *R. aerolata* ** | 140917-MRSA-09<sup>T</sup> | KU878871 | GCA_014635585.1 | 97.7 |
| 3 | *R. endophytica* | YIM 67072<sup>T</sup> | KC806052 | – | 97.6 |
| 4 | *R. terrae* ** | L-143<sup>T</sup> | DQ822568 | GCA_014705925.2 | 97.4 |
| 5 | *R. nasimurium* * | CCUG 35957<sup>T</sup> | AJ131121 | GCA_014217215.1 | 97.3 |
| 6 | *R. mucilaginosa* * | ATCC 25296<sup>T</sup> | ACVO01000020 | GCA_000175615.1 | 96.9 |
| 7 | *K. rosea* * | DSM 20447<sup>T</sup> | X87756 | GCA_006717035.1 | 96.8 |
| 8 | *K. polaris* * | CMS 76or<sup>T</sup> | JSUH01000031 | GCA_000786195.1 | 96.8 |
| 9 | *K. uropygioeca* | 257<sup>T</sup> | MF510150 | – | 96.7 |
| 10 | *K. uropygialis* | 36<sup>T</sup> | MF510149 | – | 96.7 |
| 11 | *K. subflava* * | YIM 13062<sup>T</sup> | KR610332 | GCA_012163155.1 | 96.6 |
| 12 | *R. halotolerans* * | YIM 90716<sup>T</sup> | DQ979377 | GCA_004136635.1 | 96.6 |
| 13 | *R. koreensis* * | P31<sup>T</sup> | FJ607312 | GCA_009734705.1 | 96.6 |
| 14 | *R. dentocariosa* * | ATCC 17931<sup>T</sup> | CP002280 | GCA_000164695.2 | 96.5 |
| 15 | *K. palustris* ** | DSM 11925<sup>T</sup> | Y16263 | GCA_001275345.1 | 96.5 |
| 16 | *A. crystallopoietes* * | DSM 20117<sup>T</sup> | FNKH01000002 | GCA_002849715.1 | 96.5 |
| 17 | *K. aegyptia* | YIM 70003<sup>T</sup> | DQ059617 | – | 96.4 |
| 18 | *K. salina* ** | Hv14b<sup>T</sup> | LT674162 | GCA_013368965.1 | 96.4 |
| 19 | *K. dechangensis* ** | NEAU-ST5-33<sup>T</sup> | JQ762279 | GCA_014636775.1 | 96.4 |
| 20 | *K. rhizophila* * | TA68<sup>T</sup> | RCOM01000116 | GCA_003667225.1 | 96.3 |
| 21 | *K. turfanensis* * | HO-9042<sup>T</sup> | CP014480 | GCA_001580365.1 | 96.3 |
| 22 | *K. tytonis* * | 442<sup>T</sup> | MG547562 | GCA_003226895.2 | 96.3 |
| 23 | *K. arsenatis* | CM1E1<sup>T</sup> | KM874399 | – | 96.3 |
| 24 | *K. varians* * | NBRC 15358<sup>T</sup> | BJNW01000033 | GCA_006539885.1 | 96.2 |
| 25 | *K. salsicia* * | 104<sup>T</sup> | GQ352404 | GCA_007681555.1 | 96.2 |
| 26 | *K. himachalensis* | K07-05<sup>T</sup> | AY987383 | – | 96.2 |
| 27 | *K. gwangalliensis* | SJ2<sup>T</sup> | EU286964 | – | 96.1 |
| 28 | *K. flava* * | HO-9041<sup>T</sup> | CP013254 | GCA_001482365.1 | 96.0 |
| 29 | *K. indica* * | NIO-1021<sup>T</sup> | FXAC01000036 | GCA_003667205.1 | 95.9 |
| 30 | *K. coralli* ** | SCSIO 13007<sup>T</sup> | KF895368 | GCA_008831305.1 | 95.9 |
| 31 | *K. carniphila* * | CCM 132<sup>T</sup> | AJ622907 | GCA_004136555.1 | 95.9 |
| 32 | *K. atrinae* * | P30<sup>T</sup> | FJ607311 | GCA_000286355.1 | 95.9 |
| 33 | *R. kristinae* * | NBRC 15354<sup>T</sup> | BCSM01000028 | GCA_001570865.1 | 95.8 |

Table 1 summarizes the data obtained for 33 type (^T^) strains of the genera *Rothia*, *Kocuria*, and *Arthrobacter* in the “Valid names only” category. These strains are related to Isolate SG in the 16S rRNA-based test, with values of pairwise genetic sequence identity *I* > 95%. As stated on the web page (Understanding Results, https://help.microbial-genomes.org/understanding-results#distance), *I* values of 95–98.6% correspond to strains belonging to the same genus and those of 92–95% correspond to strains belonging to the same family. According to the data of (Yarza *et al*., 2014), the cutoff threshold for assigning strains to the same genus is *I* > 94.5% and that for assigning strains to the same family is *I* > 86.5%. For assigning strains to the same species, the condition *I* > 98.6% is now accepted (Stackebrandt and Ebers, 2006; Kim *et al*., 2014), in contrast to the previously used *I* > 97% (Tindall *et al*., 2010).

A phylogenetic tree was constructed with the “Advanced” option of the NGPhylogeny web service (Lemoine *et al*., 2019; NGPhylogeny.fr, https://ngphylogeny.fr) and with the FastME 2.0 program (FastME 2.0, http://www.atgc-montpellier.fr/fastme). We used multiple sequence alignment (MSA) of the 16S rRNA genes of Isolate SG and the strains listed in Table 1.

To obtain quantitative data on the belonging of strains to a particular taxonomic category, we used the average nucleotide identity (ANI) and average amino acid identity (AAI) matrices (ANI/AAI-Matrix, http://enve-omics.ce.gatech.edu/g-matrix; Jain *et al*., 2022; Rodriguez-R and Konstantinidis, 2014; Goris *et al*., 2007) and their corresponding evolutionary distance matrices obtained from the results of whole genome sequencing of the strains’ DNA. References to the genomes of 20 strains marked with asterisks in Table 1 are given in the output data of the program (Search EzBioCloud Database, https://www.ezbiocloud.net). Additionally, on the website (Genome Information by Organism, https://www.ncbi.nlm.nih.gov/genome/browse/#!/overview), we found references to six more genomes of strains of the species listed in Table 1 and marked with a double asterisk. Thus, it became possible to take into consideration 26 whole genomes of strains across the range of *I* values from Table 1.

Tests based on ANI and AAI form the basis of the Microbial Genomes Atlas (MiGA) genomic and metagenomic data management and processing system (Rodriguez-R *et al*., 2020) for the phylogenetic classification and cataloging of microbial genomes and analysis of their gene content. We used the TypeMat version (MiGA/TypeMat/New, http://microbial-genomes.org/query_datasets/new?project_id=20), which determines the taxonomic position, novelty rank, and content of the query genome and finds the closest genomes of the type strains of officially named species from the TypeMat specialized database (Rodriguez-R *et al*., 2020) (16,345 reference genomes as of spring 2023).

In the output generated by the programs (ANI/AAI-Matrix, http://enve-omics.ce.gatech.edu/g-matrix), the phylogenetic trees built from evolutionary distance matrices do not accompany by results of branch significance calculations similar to bootstrap analysis in taxonomic studies using MSA (Lemoine et al., 2018). To address this gap, we applied the RΕQ program (REQ, https://gitlab.pasteur.fr/GIPhy/REQ), which is designed to estimate branch significance values for phylogenetic trees based on distance matrices. The closer to 100 is the quantitative value in the RΕQ output (Guénoche and Garreta, 2001), the more fully the corresponding branch is supported by the pairwise evolutionary distances.

We also determined the taxonomic position of Isolate SG on the basis of the phylogenetic system (Parks *et al*., 2017; Parks *et al*., 2018; Parks *et al*., 2020). The system is aimed at constructing the Genome Taxonomy Database (GTDB) (https://gtdb.ecogenomic.org) of bacteria and archaea with regard to monophyly and relative evolutionary divergence of species (Parks *et al*., 2018). The GTDB classification is a version of multilocus sequence analysis (MLSA) (Glaeser and Kampfer, 2015) that uses a set of concatenated amino acid sequences of ubiquitous single-copy proteins. The genes of these proteins are classified as housekeeping genes and number 120 in the case of bacteria and 122 in the case of archaea (Parks *et al*., 2017; Parks *et al*., 2018; Parks *et al*., 2020). We used the GTDB-Tk taxonomic classification toolkit (Chaumeil *et al*., 2022) in its online version (GTDB-Tk Classify, https://kbase.us/applist/apps/kb_gtdbtk/run_kb_gtdbtk).

MLSA-based methods lack their own quantitative criteria for the identification and demarcation of taxonomic ranks (Glaeser and Kampfer, 2015). However, this limitation was overcome in part in (Parks *et al*., 2020), in which the GTDB approach was supplemented by using quantitatively defined clusters of species based on ANI (ANI/AAI-Matrix, http://enve-omics.ce.gatech.edu/g-matrix; Jain *et al*., 2022; Rodriguez-R and Konstantinidis, 2014; Goris *et al*., 2007) and selected representative genomes of species. ANI values ≥ 95% combine them within species in most taxonomic studies using this test (Jain *et al*., 2018; Konstantinidis and Tiedje, 2005; Olm *et al*., 2020). As an additional tool for species identification, the alignment fraction (AF), the percentage of orthologous regions shared between the genomes under consideration, was also used in (Parks *et al*., 2020).

We note that the GTDB technology, like the aforementioned 16S rRNA-, ANI-, and AAI-based tests, remains within the paradigm of vertically inherited genotypic traits of prokaryotes, because it uses the markers from the core part of the pangenome without consideration of the effects of horizontal gene transfer (HGT) in its accessory part. The effects of HGT largely control the variety of phenotypic traits, determining, in particular, the ability of bacteria and archaea to adapt and function in diverse and frequently changing ecological niches (Koonin, 2012). These traits are taken into account when the systematic position of objects is evaluated on the basis of the polyphasic approach, which is very common in the traditional systematics of prokaryotes (Oren and Garrity, 2014). Hence follows the obvious conventionality of phylogenetic analysis within any scheme using only vertically inherited phylogenetic markers, as does possible disagreement of its results with the traditional prokaryote classification and nomenclature (Shchyogolev, 2021). This probably explains, in particular, the need to verify prokaryote classification and nomenclature with the corresponding classification changes, which turned out to be relevant for about 60% of the GTDB objects (Genome Taxonomy Database, https://gtdb.ecogenomic.org) analyzed in (Parks *et al*., 2018).

The systematic position of Isolate SG at the whole-genome level with respect to the objects from the UniProt protein structure database (UniProtKB, https://www.uniprot.org) (250 million records) was also evaluated by us with the AAI-profiler program (AAI-profiler, http://ekhidna2.biocenter.helsinki.fi/AAI; Medlar *et al*., 2018). As the input data, AAI-profiler uses the proteome of the strain under study, determining the AAI between the query and the species members in UniProt and the coverage, i.e., the proportion of matched protein pairs (matched fraction, MF). In particular, the program identifies and visualizes (1) possible contradictions in the classification of pro- and eukaryotes and (2) microbial contamination (Medlar *et al*., 2018).

To visualize and analyze phylogenetic structures with relatively small numbers of operational taxonomic units (OTUs) (tens and hundreds), we used the MEGA11 program (MEGA, https://www.megasoftware.net). For trees with hundreds of thousands of OTUs, we used the Archaeopteryx program (Archaeopteryx. https://sites.google.com/site/cmzmasek/christian-zmasek/software/archaeopteryx?authuser=0).

Table 2 summarizes the phylogenetic tests used, with quantitative criteria for taxon demarcation.

**Table 2.** Phylogenetic tests used.

| Test | Description and reference | Use and reference |
| --- | --- | --- |
| 16S rRNA | Comparison of the gene sequences of small subunit ribosomal RNA (Search EzBioCloud Database, https://www.ezbiocloud.net). | Demarcation of objects on the basis of the sequence identity $I$ within:<br>○ the species ($I > 98.6\%$)<br>○ the genus ($I > 95\%$)<br>○ the family ($I > 87\text{--}92\%$) |
| ANI | Estimation of the evolutionary distances between genomes on the basis of the average nucleotide identity (ANI/AAI-Matrix, http://enve-omics.ce.gatech.edu/g-matrix). | Demarcation of objects on the basis of the parameter values within:<br>○ the species (ANI $\geq 95\%$)<br>○ the genus (ANI $\geq 80\%$)<br>Intergeneric identification at AAI $< 80\%$ is fundamentally limited by the capabilities of the method |
| AAI | Estimation of the evolutionary distances between proteomes on the basis of the average amino acid identity (ANI/AAI-Matrix, http://enve-omics.ce.gatech.edu/g-matrix). | Demarcation of objects on the basis of the parameter values within:<br>○ the species (AAI $\geq 90\%$)<br>○ the genus (ANI $\geq 65\%$)<br>○ the family (ANI $> 45\%$) |
| TypeMat | Estimation of the evolutionary distances between the query genome and the reference genomes in the TypeMat database by using AAI and ANI (MiGA/TypeMat/New, http://microbial-genomes.org/query_datasets/new?project_id=20). | Determination of the taxonomic classification and novelty rank of the genome and evaluation of its contamination with possible HGT (Rodriguez-R *et al.*, 2020). |
| GTDB-Tk | MLSA with pooling of the amino acid sequences of ubiquitous single-copy proteins of prokaryotes, with quantitative identification of species clusters by ANI $> 95\%$, and with account taken of the percentage of orthologous regions (AF) (GTDB-Tk, https://kbase.us/applist/apps/kb_gtdbtk/run_kb_gtdbtk). | Assignment of taxonomic classifications to bacterial and archaeal genomes within the GTDB (Genome Taxonomy Database, https://gtdb.ecogenomic.org) |
| AAI-profiler | Determination of the AAI between query proteomes and species members in the UniProt database and of the coverage (MF) (AAI-profiler, http://ekhidna2.biocenter.helsinki.fi/AAI). | Evaluation of the systematic position of the strain relative to the UniProt database objects, with account taken of the AAI threshold values for taxa demarcation. |

## 3. Results and Discussion

The *I* value (Table 1, red color) indicates that Isolate SG and *R. amarae* strain JCM 11375 belong to the same species. The > 95% pairwise *I* values, obtained for Isolate SG with all strains (Table 1, column 6), are the basis for assigning Isolate SG to the same genus with each of the indicated 33 type strains (16S rRNA-based test). However, within the traditional taxonomy, these strains have been described by their authors as members of three genera: *Rothia*, *Kocuria*, and *Arthrobacter*. This illustrates the conventionality of the traditional prokaryote classification and nomenclature and the need for their essential revision and clarification. This need has been clearly identified in recent years on the basis of phylogenetic studies using whole genome data (Parks *et al*., 2017; Parks *et al*., 2018; Parks *et al*., 2020; Genome Taxonomy Database, https://gtdb.ecogenomic.org).

Fig. 1 shows the phylogram of 16S rRNA gene sequences obtained with FastME 2.0 (FastME 2.0, http://www.atgc-montpellier.fr/fastme) for the strains listed in Table 1. The red dots mark the nodes of four clusters (monophyletic groups a–d) comprising sets of 5 to 12 *Rothia*, *Kocuria*, and *Arthrobacter* strains. As it follows from the results of determination of the percentage identity matrix (data not shown), the pairwise values of *I* between strains within these monophyletic groups mostly satisfy the condition *I* > 95%. This indicates that these strains belong to the same genus within 16S rRNA technology according to the criteria (Understanding Results, https://help.microbial-genomes.org/understanding-results#distance; Yarza *et al*., 2014; Stackebrandt and Ebers, 2006; Kim *et al*., 2014). Meanwhile, between clusters, *I* values generally satisfy the 87–92% < *I* < 95% condition, which demarcates genera within a family. However, both within and outside the clusters, there are a notable number of deviations of *I* values from these quantitative criteria of taxon demarcation. This fact illustrates their conventionality as averaged characteristics within fairly broad distributions of the corresponding statistical data with 16S rRNA sequences (Luo et al., 2014).

**Fig. 1.**
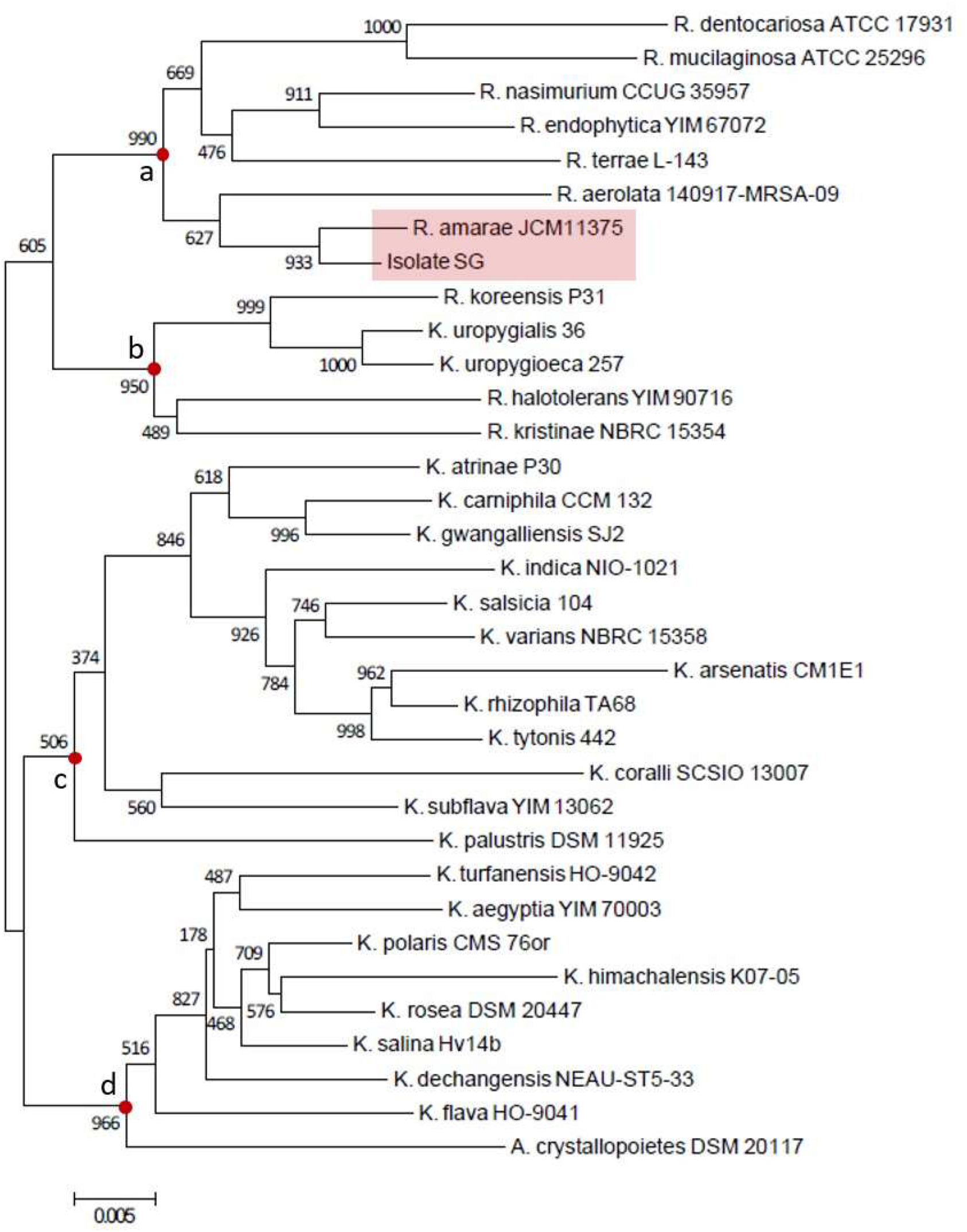
Phylogram (FastME) of the 16S rRNA gene sequences of the *Rothia*, *Kocuria*, and *Arthrobacter* type strains related to Isolate SG (Table 1). The numbers near the nodes indicate statistical support for the branches (absolute units) in 1000 bootstrapping cycles.

More unambiguous taxonomic estimates can be expected from the use of the ANI and AAI matrices on the basis of the results of whole-genome DNA sequencing of the strains (ANI/AAI-Matrix, http://enve-omics.ce.gatech.edu/g-matrix; Jain *et al*., 2018; Rodriguez-R and Konstantinidis, 2014; Goris *et al*., 2007). In Table 1, GCA_007666515.1 was initially specified as the genome corresponding to *R. amarae* strain JCM 11375^T^, which is the most closely related to Isolate SG. The application of the GTDB-Tk software package (GTDB-Tk, https://kbase.us/applist/apps/kb_gtdbtk/run_kb_gtdbtk) resulted in the following taxonomic classification of this genome: d Bacteria;p Actinobacteriota;c Actinomycetia; o Actinomycetales;f Micrococcaceae;g Rothia;s Rothia sp001683935, with the reference genome GCF_001683935.1 of *Rothia* sp. strain ND6WE1A. The resulting pairwise value of ANI = 98.87% and the value of AF = 96% indicate high intraspecies relatedness between *R. amarae* JCM 11375^T^ and *Rothia* sp. ND6WE1A. The note states that “topological placement and ANI have congruent species assignments.”

Note that the GCA_007666515.1 genome is described in the NCBI database record as “anomalous assembly” owing to the “unverified source organism.” This was in no way evident when the genome was processed with GTDB-Tk (GTDB-Tk, https://kbase.us/applist/apps/kb_gtdbtk/run_kb_gtdbtk) to determine its most probable taxonomic classification and clarify the species identity of the strain under study on the basis of ANI. Nonetheless, in subsequent whole-genome tests, we used as a representative of Isolate SG the GCF_001683935.1 genome⸻the reference genome (ANI = 98.87%) for GCA_007666515.1 in the GTDB (Genome Taxonomy Database, https://gtdb.ecogenomic.org).

Fig. 2 shows the phylogram of the ANI of the 26 strains being considered. The phylogram is presented in the output data of the software package (ANI/AAI-Matrix, http://enve-omics.ce.gatech.edu/g-matrix) and was constructed by the BioNJ method by using a matrix of ANI values. The red dots mark the nodes of five monophyletic groups (a–e) comprising sets of three to seven *Rothia*, *Kocuria*, and *Arthrobacter* strains. The results reflected in the matrix of pairwise ANI values (data not shown) showed that in several strains, there were almost no groups of genes orthologous to the genes of *Rothia* sp. ND6WE1A, the reference strain for Isolate SG.

**Fig. 2.**
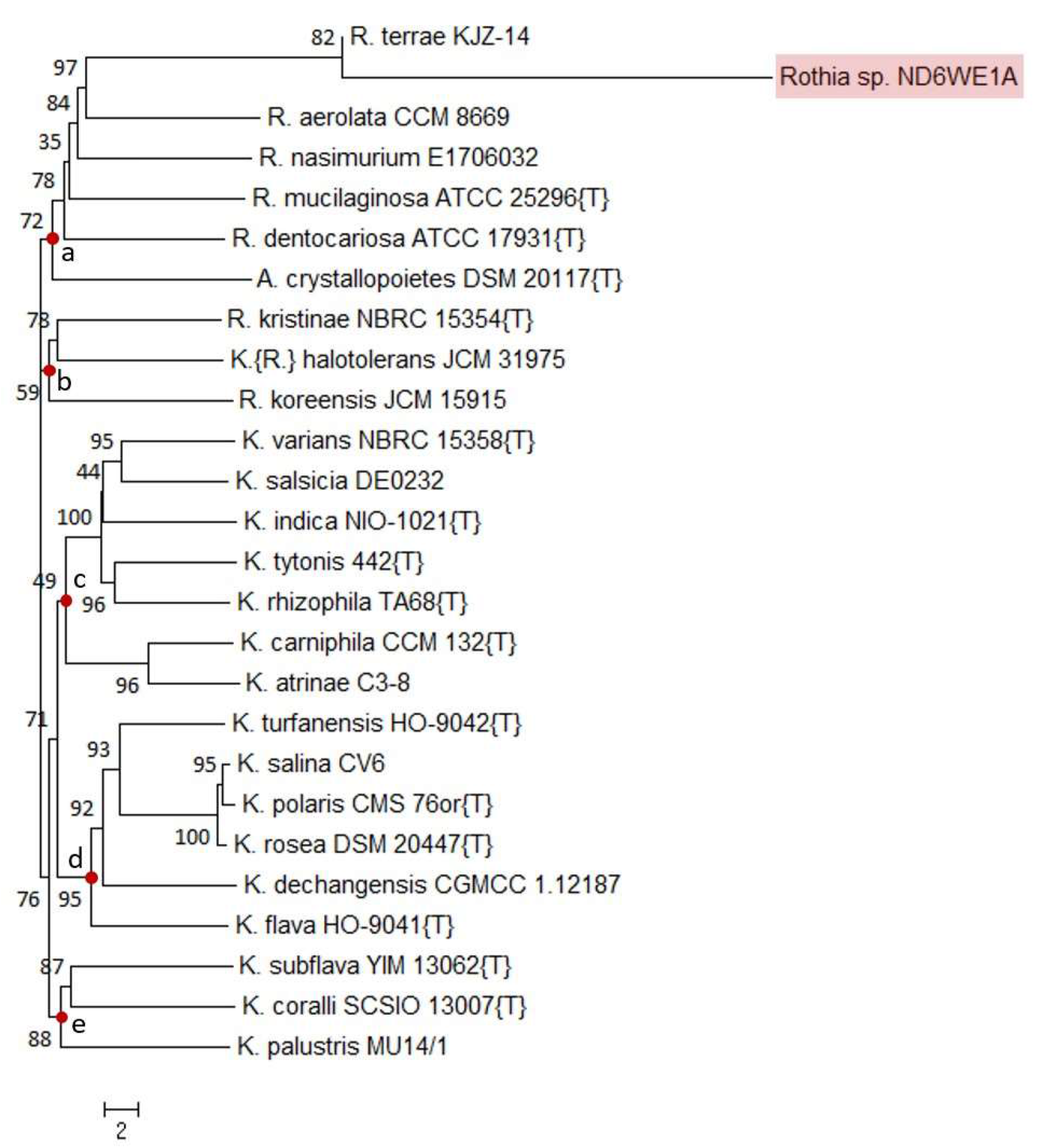
Phylogram obtained by the BioNJ method by using the distance matrix in the ANI-based test with the genomes of the strains listed in Table 1. The numbers near the nodes are branch significance estimates made with REQ (https://gitlab.pasteur.fr/GIPhy/REQ).

This explains the greatest evolutionary distance for *Rothia* sp. ND6WE1A among all the 26 strains taken into consideration (Fig. 2).

As in the case of 16S rRNA, the matrix of pairwise ANI values did not ensure unambiguous taxonomic demarcation of the strains under study above the genus level (ANI < 80%), both within and between the monophyletic groups in Fig. 2. This is associated with the limited sensitivity of the ANI-based test at ANI values ≤ 80% (Rodriguez-R and Konstantinidis, 2014). For more reliable quantitative demarcation of strains at the family level, the developers of the resource (ANI/AAI-Matrix, http://enve-omics.ce.gatech.edu/g-matrix) recommended the use of the AAI-based test.

Following this recommendation, we took advantage of the software package (ANI/AAI-Matrix, http://enve-omics.ce.gatech.edu/g-matrix) to obtain a matrix of AAI values and the corresponding phylogenetic constructs by using the proteomes of the same 26 *Rothia*, *Kocuria*, and *Arthrobacter* strains. In the resulting phylogram (Fig. 3), the red dots indicate the nodes of 5 monophyletic groups a–e, comprising 2 to 13 strains. The results of determining the matrix of pairwise AAI values (Fig. 4) show a fairly clear distribution of strains between these five groups, marked with dashed rectangles in Fig. 4, with intrageneric (AAI ≥ 65%, red) and intraspecies (AAI > 90%, yellow) AAI values (Understanding Results. https://help.microbial-genomes.org/understanding-results#distance). To demarcate genera at the family level, the condition 45% < AAI < 65% (colorless cells) is given on the web page (Understanding Results. https://help.microbial-genomes.org/understanding-results#distance). These inequalities are based on averaged characteristics and are generally consistent with the statistical analysis results given in (Luo *et al*., 2014). Note the higher branch support in the AAI-based test (Fig. 3), as compared with that in the ANI-(Fig. 2) and 16S rRNA-based (Fig. 1) tests.

**Fig. 3.**
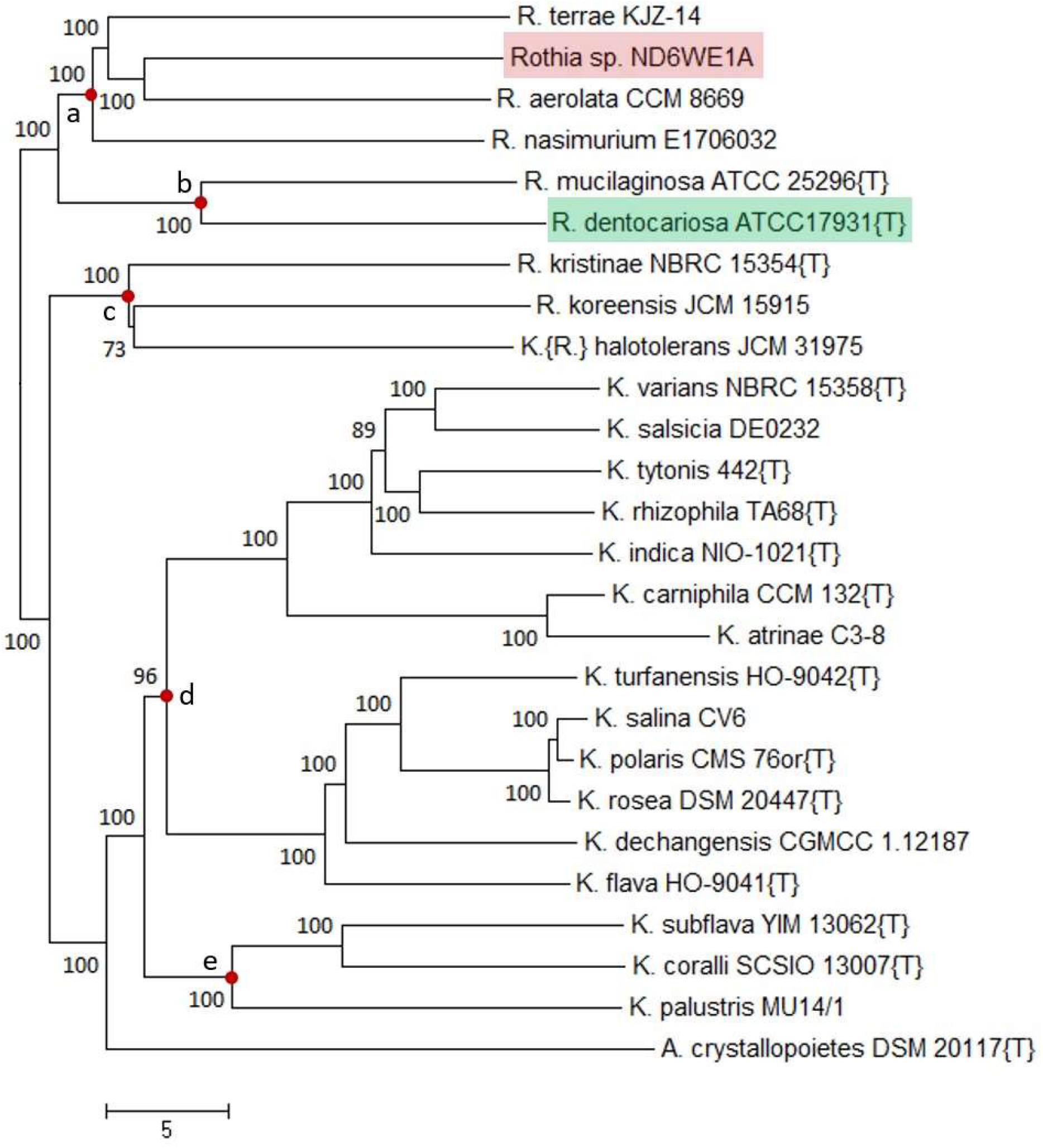
Phylogram obtained by the BioNJ method by using the distance matrix in the AAI-based test with the genomes of the strains listed in Table 1. The numbers near the nodes are branch significance estimates made with REQ (https://gitlab.pasteur.fr/GIPhy/REQ).

**Fig. 4.**
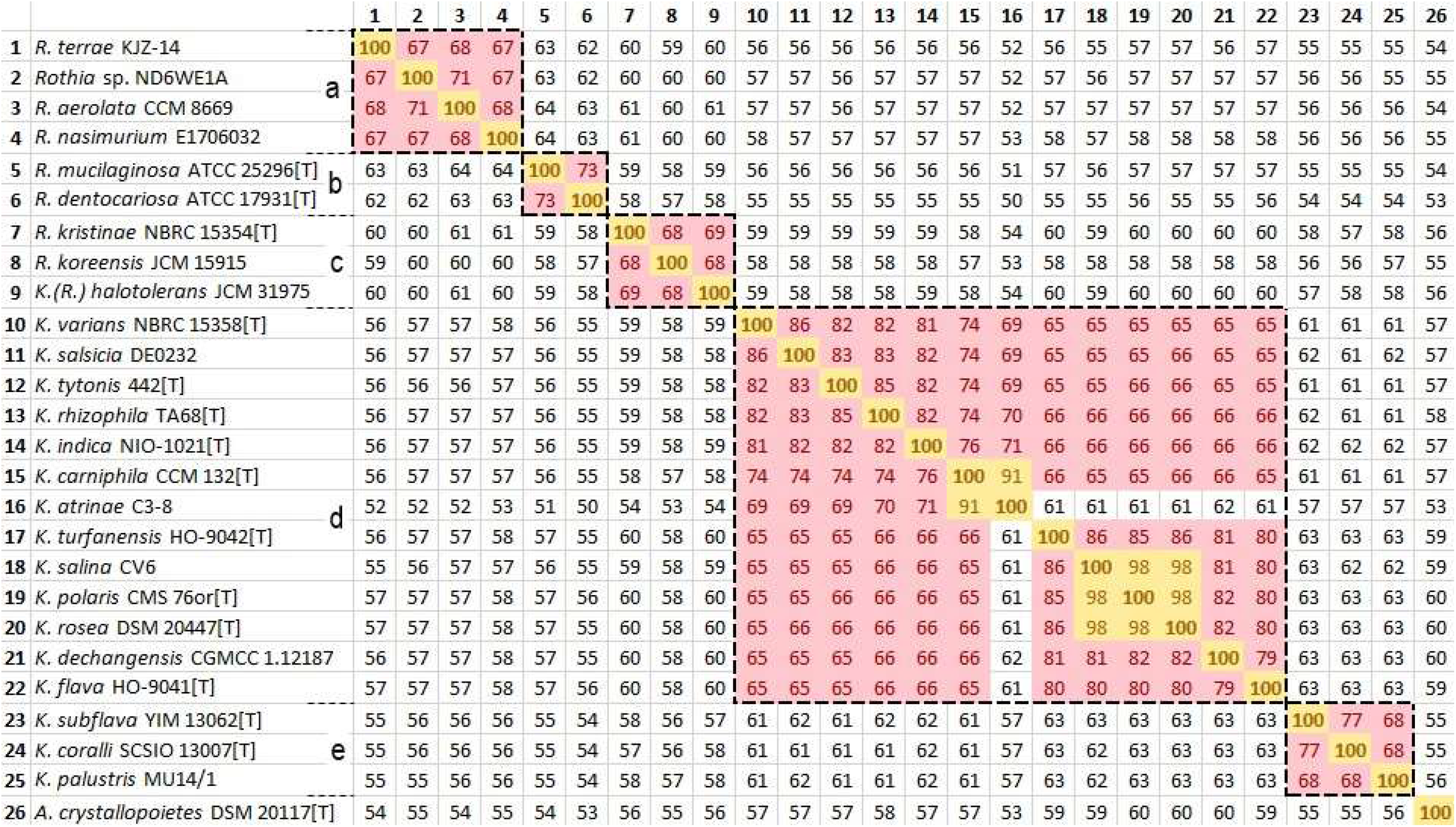
AAI value matrix obtained from the distance matrix presented in the output data of the software package (ANI/AAI-Matrix, http://enve-omics.ce.gatech.edu/g-matrix). The dashed rectangles mark the fragments corresponding to monophyletic groups a–e, whose nodes are marked with red dots in Fig. 3.

Table 3 presents the distribution of the strains by monophyletic group, as found by phylogenetic analysis using the tests based on 16S rRNA, ANI, and AAI (Figs. 1–4). We note the general qualitative agreement of the clustering of the strains for the three tests. However, only the AAI-based test provided a sufficiently consistent association of strains within the same genus at the quantitative level, which agrees with the general recommendations found on the web page (Understanding Results, https://help.microbial-genomes.org/understanding-results#distance) and in the publications (Rodriguez-R and Konstantinidis, 2014; Goris *et al*., 2007).

**Table 3.** Clustering of the strains related to Isolate SG in 16S rRNA-based test.

| Test |  |  |
| --- | --- | --- |
| 16S rRNA | ANI | AAI |
| *R. dentocariosa* ATCC 17931<sup>T</sup><br>*R. mucilaginoso* ATCC 25296<sup>T</sup><br>*R. nasimurium* CCUG 35957<sup>T</sup><br>*R. endophytica* YIM 67072<sup>T</sup><br>*R. terrae* L-143<sup>T</sup><br>*R. aerolata* 140917-MRSA-09<sup>T</sup><br>*R. amarae* JCM 11375<sup>T</sup><br>**Isolate SG** | *R. dentocariosa* ATCC 17931<sup>T</sup><br>*R. mucilaginoso* ATCC 25296<sup>T</sup><br>*R. nasimurium* E1706032<br>*R. terrae* KJZ-14<br>*R. aerolata* CCM 8669<br>*A. crystallopoietes* DSM 20117<sup>T</sup><br>*Rothia* sp. ND6WE1A (representative of **Isolate SG**) | *R. dentocariosa* ATCC 17931<sup>T</sup><br>*R. mucilaginoso* ATCC 25296<sup>T</sup><br>*R. nasimurium* E1706032<br>*R. terrae* KJZ-14<br>*R. aerolata* CCM 8669<br>*Rothia* sp. ND6WE1A (representative of **Isolate SG**) |
| *R. koreensis* P31<sup>T</sup><br>*K. uropygialis* 36<sup>T</sup><br>*K. uropygioeca* 257<sup>T</sup><br>*R. halotolerans* YIM 90716<sup>T</sup><br>*R. kristinae* NBRC 15354<sup>T</sup> | *R. koreensis* JCM 15915<br>*K.(R.) halotolerans* JCM 31975<br>*R. kristinae* NBRC 15354<sup>T</sup> | *R. koreensis* JCM 15915<br>*K.(R.) halotolerans* JCM 31975<br>*R. kristinae* NBRC 15354<sup>T</sup> |
| *K. atrinae* P30<sup>T</sup><br>*K. carniphila* CCM 132<sup>T</sup><br>*K. gwangalliensis* SJ2<sup>T</sup><br>*K. indica* NIO-1021<sup>T</sup><br>*K. salsicia* 104<sup>T</sup><br>*K. varians* NBRC 15358<sup>T</sup><br>*K. arsenatis* CM1E1<sup>T</sup><br>*K. rhizophila* TA68<sup>T</sup><br>*K. tytonis* 442<sup>T</sup><br>*K. coralli* SCSIO 13007<sup>T</sup><br>*K. subflava* YIM 13062<sup>T</sup><br>*K. palustris* DSM 11925<sup>T</sup> | *K. atrinae* C3-8<br>*K. carniphila* CCM 132<sup>T</sup><br>*K. indica* NIO-1021<sup>T</sup><br>*K. salsicia* DE0232<br>*K. varians* NBRC 15358<sup>T</sup><br>*K. rhizophila* TA68<sup>T</sup><br>*K. tytonis* 442<sup>T</sup><br>*K. turfanensis* HO-9042<sup>T</sup><br>*K. polaris* CMS 76or<sup>T</sup><br>*K. rosea* DSM 20447<sup>T</sup><br>*K. salina* CV6<br>*K. dechangensis* CGMCC 1.12187<br>*K. flava* HO-9041<sup>T</sup> | *K. atrinae* C3-8<br>*K. carniphila* CCM 132<sup>T</sup><br>*K. indica* NIO-1021<sup>T</sup><br>*K. salsicia* DE0232<br>*K. varians* NBRC 15358<sup>T</sup><br>*K. rhizophila* TA68<sup>T</sup><br>*K. tytonis* 442<sup>T</sup><br>*K. turfanensis* HO-9042<sup>T</sup><br>*K. polaris* CMS 76or<sup>T</sup><br>*K. rosea* DSM 20447<sup>T</sup><br>*K. salina* CV6<br>*K. dechangensis* CGMCC 1.12187<br>*K. flava* HO-9041<sup>T</sup> |
| *K. turfanensis* HO-9042<sup>T</sup><br>*K. aegyptia* YIM 70003<sup>T</sup><br>*K. polaris* CMS 76or<sup>T</sup><br>*K. himachalensis* K07-05<sup>T</sup><br>*K. rosea* DSM 20447<sup>T</sup><br>*K. salina* Hv14b<sup>T</sup><br>*K. dechangensis* NEAU-ST5-33<sup>T</sup><br>*K. flava* HO-9041<sup>T</sup><br>*A. crystallopoietes* DSM 20117<sup>T</sup> | *K. coralli* SCSIO 13007<sup>T</sup><br>*K. subflava* YIM 13062<sup>T</sup><br>*K. palustris* MU14/1 | *K. coralli* SCSIO 13007<sup>T</sup><br>*K. subflava* YIM 13062<sup>T</sup><br>*K. palustris* MU14/1<br>*A. crystallopoietes* DSM 20117<sup>T</sup> |

In this case, the strains collected within separate monophyletic group a–e in Figs. 3 and 4 should be considered belonging to the same genus (AAI ≥ 65%). By the AAI < 65% criterion, however, the evolutionary distances between these groups correspond to the intergeneric level within the family *Micrococcaceae*. Consequently, the genus affiliation of the strains in monophyletic groups a, b, and c, originally assigned by their authors to the genus *Rothia*, and d and e, originally assigned to the genus *Kocuria*, requires clarification (with possible renaming). *Rothia* sp. ND6WE1A, the reference strain for Isolate SG at the whole genome level, and *R. dentocariosa* ATCC 17931^T^, the type species for the genus *Rothia*, are part of groups a and b (Fig. 3, 4), separated by intergeneric AAI values. In the AAI-based test, *A. crystallopoietes* strain DSM 20117^T^ and partly *R. mucilaginosa* ATCC 25296^T^ are found outside the genera *Rothia* and *Kocuria*; in the 16S rRNA-based test (Table 1, column 5), these strains show an intrageneric level of sequence identity to Isolate SG *I* > 95%.

These observations on *Rothia* sp. ND6WE1A are consistent with the results of MiGA technology (Rodriguez-R *et al*., 2020; MiGA/TypeMat/New, http://microbial-genomes.org/query_datasets/new?project_id=20). These results indicate that this genome most probably belongs to the order *Micrococcales* (*p* = 0.0024) and probably belongs to the genus *Rothia* (*p* =0.31). However, the “Taxonomic novelty” section states that it most probably belongs to a species not included in the TypMat database (MiGA/TypeMat/New, http://microbial-genomes.org/query_datasets/new?project_id=20) (*p* = 0.0021), which comprises genomes of type strains (including *Rothia dentocariosa* ATCC 17931^T^), the highest taxonomic rank with *p* ≤ 0.01. The data presented in MyTaxa Scan testify to the detection in the genome of *Rothia* sp. ND6WE1A of regions with an unusual taxonomic distribution. These regions are shown in the MyTaxa Scan diagram (not presented). According to the publications (Rodriguez-R *et al*., 2020; Luo *et al*., 2014; Rodriguez-R *et al*., 2018), they can be interpreted as the result of HGT, because the estimates presented in the “Quality (essential genes)” section show zero contamination of the core part of the genome.

*Rothia* sp. ND6WE1A, a strain representative of Isolate SG, is part of the GTDB R06-RS202 reference tree, included in the file bac120_r202.tree, with 254,091 OTUs (GTDB Data, https://data.gtdb.ecogenomic.org/releases/release202/202.0/). Fig. 5 shows a *Micrococcaceae* subtree fragment that is a monophyletic group comprising strains assigned to the genus *Rothia* in GTDB R06-RS202. This fragment was extracted by us from the file bac120_r202.tree with the program (Archaeopteryx, https://sites.google.com/site/cmzmasek/christian-zmasek/software/ archaeopteryx?authuser=0). In the phylogenetic tree of Fig. 5, including 22 OTUs, we replaced the genome designations in the file bac120_r202.tree (GTDB Data, https://data.gtdb.ecogenomic.org/releases/release202/202.0/) [access codes to the NCBI genome assemblies (Genome Information by Organism. https://www.ncbi.nlm.nih.gov/genome/browse/#!/overview)] by the strain names given in the corresponding entries in the NCBI database. The red dots denote cluster nodes⸻three isolated monophyletic groups a–c within the g_*Rothia* fragment of the file bac120_r202.tree (GTDB Data, https://data.gtdb.ecogenomic.org/releases/ release202/202.0/).

**Fig. 5.**
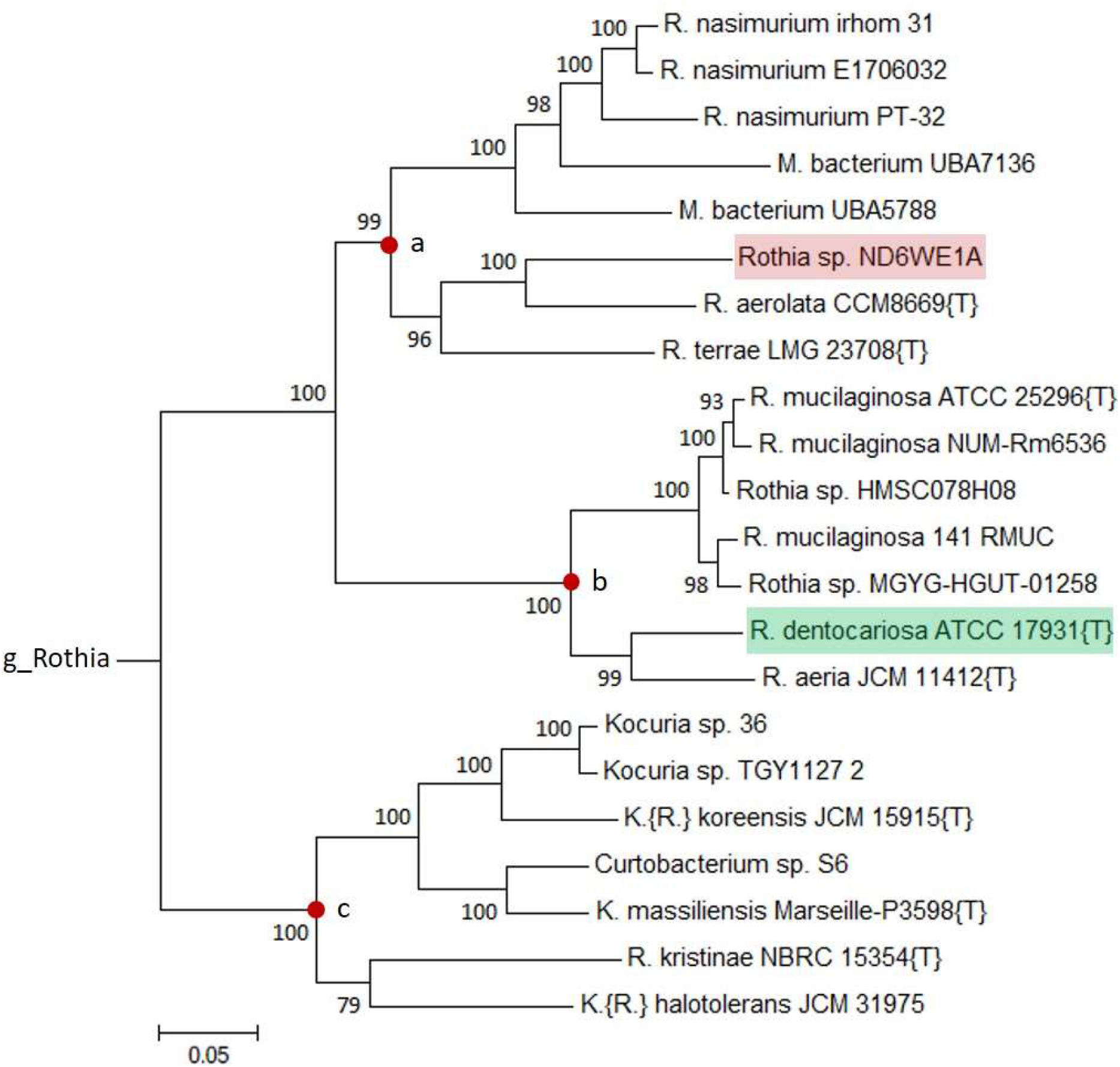
Fragment of the reference phylogenetic tree bac120_r202.tree (GTDB Data, https://data.gtdb.ecogenomic.org/releases/release202/202.0/) for the monophyletic group assigned to the genus *Rothia* in GTDB R06-RS202.

As an example of the abovementioned need for reclassification of the taxonomic affiliations detected by GTDB technology (Parks *et al*., 2018), we note that the objects attributed by their authors to the genera *Curtobacterium* and *Kocuria* are found in the monophyletic group of *Rothia* (Fig. 5 c). A positive moment here is the elimination of the uncertainty of the systematic position of the metagenomic objects *Micrococcaceae bacterium* UBA7136 (rat intestinal metagenome) and *M. bacterium* UBA5788 (urban ground metagenome), having the status “unclassified Micrococcaceae” and found as part of one cluster in the monophyletic group of *Rothia* (Fig. 5 a). To quantify the generic assignment of the strains and metagenomic objects belonging to monophyletic groups a–c of the Fig. 5 phylogram, whose nodes are marked with red dots, we used the AAI test (ANI/AAI-Matrix, http://enve-omics.ce.gatech.edu/g-matrix) as a reliable means to distinguish between strains within the family (Rodriguez-R and Konstantinidis, 2014; Goris *et al*., 2007; Medlar *et al*., 2018). Among the 22 strains in Fig. 5, whose genomes are used in GTDB R06-RS202 (GTDB Data, https://data.gtdb.ecogenomic.org/releases/release202/202.0/), we did not find the proteomes of *M. bacterium* UBA7136, *M. bacterium* UBA5788, and *Rothia* sp. MGYG-HGUT-01258 in the NCBI database (Genome Information by Organism. https://www.ncbi.nlm.nih.gov/genome/browse/#!/overview). The results we obtained for the remaining 19 strains are shown in Figs. 6 and 7.

**Fig. 6.**
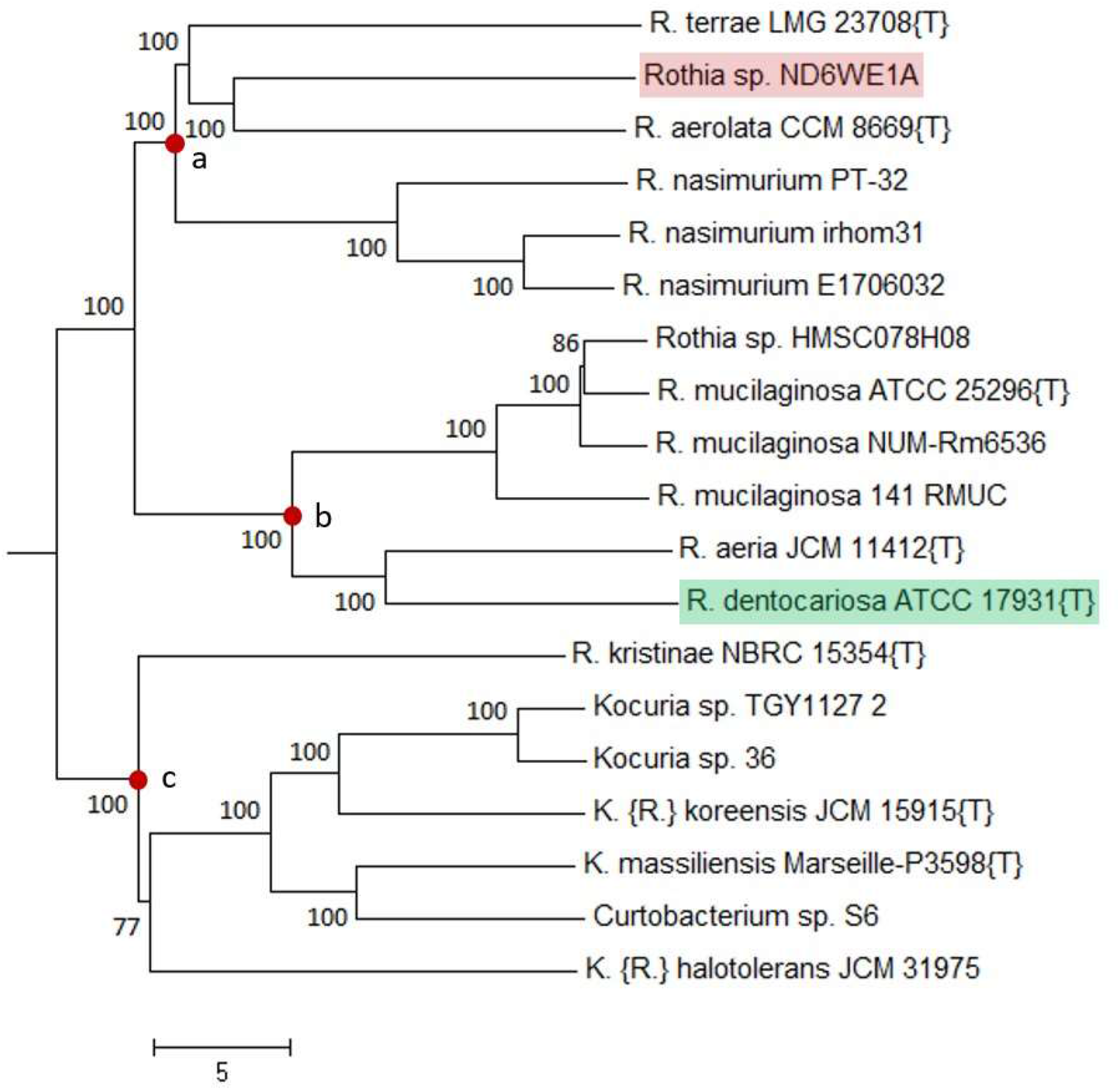
Phylogram obtained by the BioNJ method by using the distance matrix in the AAI-based test with the proteomes of the strains from the monophyletic group assigned to the genus *Rothia* in GTDB R06-RS202 (Fig. 5). The numbers near the nodes are branch significance estimates made with REQ (https://gitlab.pasteur.fr/GIPhy/REQ).

**Fig. 7.**
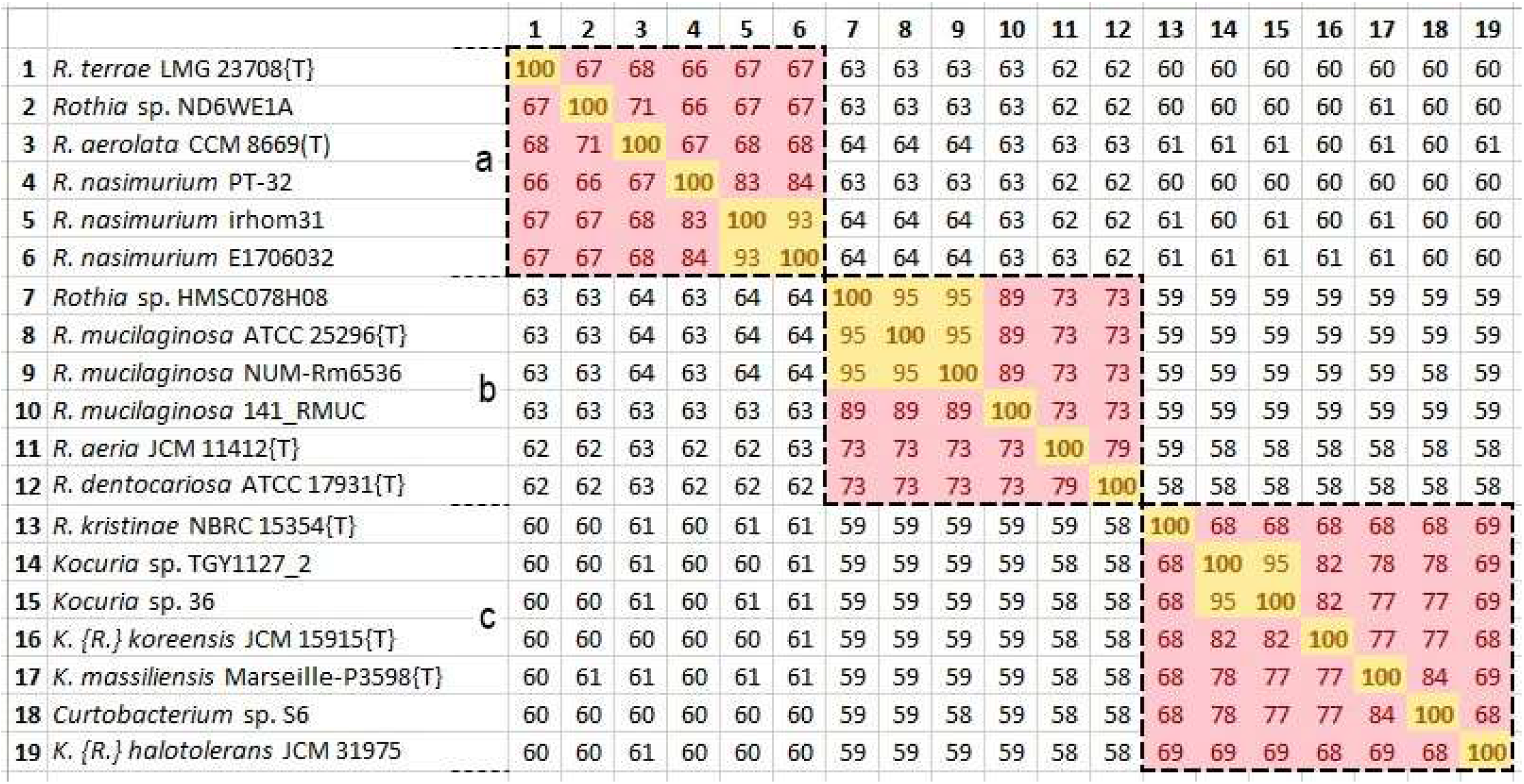
AAI value matrix obtained from the distance matrix presented in the output data of the software package (ANI/AAI-Matrix, http://enve-omics.ce.gatech.edu/g-matrix). The dashed rectangles mark the fragments corresponding to monophyletic groups a–c, whose nodes are marked with red dots in Fig. 6.

The red dots in Fig. 6 mark the nodes of three clusters corresponding to those in Fig. 5, with the same designations (a–c). The species (strain-level) and genus (cluster-level) demarcation of the OTUs in Figs. 5 and 6 is quantitatively ensured by their comparison with the matrix of AAI values in Fig. 7, in which the areas corresponding to clusters a–c in Figs. 5 and 6 are marked with dashed lines. The coloring shows cells satisfying the AAI ≥ 90% (yellow, same species) and AAI ≥ 65% (red, same genus) conditions (Understanding Results, https://help.microbial-genomes.org/understanding-results#distance; Luo *et al*., 2014). The colorless cells correspond to intergeneric pairwise values of 45% < AAI < 65%, joining clusters a–c together (Figs. 5 and 6) within the same family (Understanding Results, https://help.microbial-genomes.org/understanding-results#distance; Luo *et al*., 2014). The belonging to the same species in the AAI-based test (Fig. 7, yellow cells) is confirmed for the strains *R. nasimurium* irhom31/*R. nasimurium* E1706032; *Rothia* sp. HMSC078H08, with the pair *R. mucilaginosa* ATCC 25296^T^/*R. mucilaginosa* NUM-Rm6536; *Kocuria* sp. TGY1127_2/*Kocuria* sp. 36.

The most substantial result of using the AAI-based test in the analysis of the phylograms in Figs. 5 and 6 for the strains assigned to *Rothia* in GTDB R06-RS202 (GTDB Data, https://data.gtdb.ecogenomic.org/releases/release202/202.0/) is the < 65% intergeneric level of AAI values (Understanding Results, https://help.microbial-genomes.org/understanding-results#distance; Luo *et al*., 2014) for clusters a, b, and c (Fig. 7). In accordance with the conditions (Understanding Results, https://help.microbial-genomes.org/understanding-results#distance; Luo *et al*., 2014), these clusters should be assigned to different genera within the same family (AAI > 45%), with reclassification of the objects included in them.

Fig. 8 shows the AAI distribution diagram for the proteome of the reference strain *Rothia* sp. ND6WE1A, used as a query in the AAI-profiler test (AAI-profiler, http://ekhidna2.biocenter.helsinki.fi/AAI; Medlar *et al*., 2018). On the horizontal axis are plotted AAI values between the query and the species members in the UniProt database (UniProtKB, https://www.uniprot.org), whereas the vertical axis shows the matched fraction (MF, coverage). The icons in the diagram correspond to the species that have scored the highest marks with allowance for AAI and MF, i.e., the sum of the sequence identity values for all query proteins with established matches. Related species, grouped and colored on the basis of genus, form a characteristic “cloud” in the diagram, with AAI values reflecting the evolutionary proximity of the UniProt strains to the query strain. The horizontal axis has icons for the species for which DNA sequencing results have been obtained for individual proteins only. The icons are colored according to genus (bacteria) or order (eukaryotes). Eukaryotic species are marked with rhombuses; bacteria, with circles; archaea, with crosses; and everything else (viruses, metagenomes, unclassified samples), with squares.

**Fig. 8.**
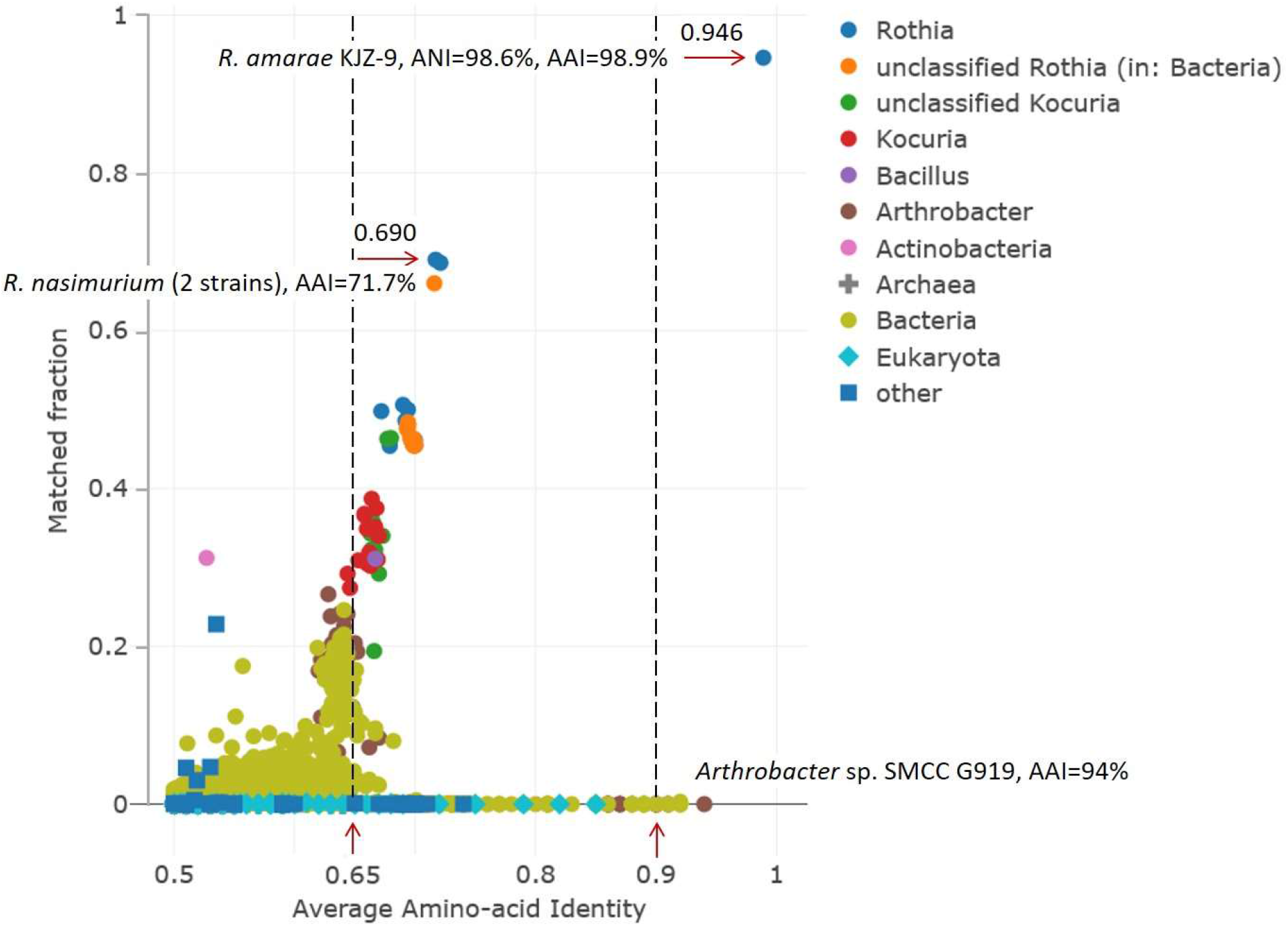
AAI distribution diagram for *Rothia* sp. ND6WE1A, as found in the AAI-profiler output data. Explanations are in the text.

The vertical dashed lines in Fig. 8 correspond to the AAI threshold values for grouping strains into genera (AAI > 0.65) and species (AAI > 0.9) according to the conditions specified on the web page (Understanding Results, https://help.microbial-genomes.org/understanding-results#distance). The strain most closely related to *Rothia* sp. ND6WE1A is *R. amarae* KJZ-9, with AAI = 98.9%, ANI = 98.6%, and MF = 0.946. Thus, all three genomes [GCA_007666515.1, *R. amarae* (*mucilaginosa*) DE0531 (originally listed in Table 1 as corresponding to *R. amarae* JCM 11375^T^ but excluded from our consideration; see above); GCA_001683935.1, *Rothia* sp. ND6WE1A; and GCA_014705945.2, *R. amarae* KJZ-9] represent the same species with high inter-strain evolutionary relatedness, characterized by ANI and AAI values of about 99%.

Another strain closely related to *Rothia* sp. ND6WE1A [AAI = 94%, which exceeds the threshold value for grouping strains at the species level (AAI > 0.9)] is *Arthrobacter* sp. SMCC G919, whose icon is located on the horizontal axis of the diagram (Fig. 8). Then come members of *Thermocrinis*, *Planobispora*, *Mycobacterium*, *Glutamicibacter*, *Micrococcus*, and other genera, which too are located on the horizontal axis in descending order of AAI values ≍ 0.9. The appearance in this diagram, grouping mostly *Micrococcaceae* members, of “noncongruent” species with a small (almost zero) MF may be associated with HGT (Medlar *et al*., 2018; Treangen and Rocha, 2011).

On the other hand, *Arthrobacter* occupies a certain position in the main part of the diagram with an MF in the range 0.03–0.25 and AAI = 62–67% in the region of 60–80%, which groups strains at the genus level (Luo *et al*., 2014). Members of *Rothia* and *Kocuria* also fall into the same area in the diagram (Fig. 8), with MFs ranging from 0.28 to 0.95. Such “incorporation” of *Kocuria* and *Arthrobacter* members (and other bacterial species/strains marked with pink, green, and light green dots in Fig. 8) into the main cluster of *Rothia*, comprising species related to the query strain *Rothia* sp. ND6WE1A, should be considered a basis for taxonomic reclassification of these members (Medlar *et al*., 2018), which was also shown above with the tests based on 16S rRNA, ANI, and AAI.

As found with AAI-profiler, the strain most closely related to *Rothia* sp. ND6WE1A (after *R. amarae* KJZ-9) is *R. nasimurium* PT-32 (Fig. 8; AAI = 71.7%, MF = 0.69). These data show a noticeable gap between *Rothia* sp. ND6WE1A (and its closely related *R. amarae* KJZ-9) and the main group of species in Fig. 8 (MF, 25% and greater), which indicates that its genome differs strongly from those of the related species, found with AAI-profiler in the UniProt database. This observation is in harmony with the results of phylogenetic studies using ANI (Fig. 2), which showed a considerable evolutionary deviation of *Rothia* sp. ND6WE1A from the 26 *Rothia*, *Kocuria*, and *Arthrobacter* strains considered.

## 4. Conclusions

The AAI-based test results (Figs. 3 and 4) indicate that *Rothia* sp. ND6WE1A, which represents Isolate SG [recovered from cells of an *A. thaliana* (L.) Heynh suspension culture (Sokolov *et al*., 2021)] at the level of the whole genomes studied, belongs to the genus *Rothia* and is part of one monophyletic group with *R. aerolata* CCM 8669 (AAI = 71%), *R. terrae* KJZ-14 (AAI = 67%), and *R. nasimurium* E1706032 (AAI = 67%) in cluster a (Figs. 3 and 4). Members of these species are also included in clusters with Isolate SG in the 16S rRNA- and ANI-based tests (Table 3).

Their sources are highly diverse in type and geography. *Rothia* sp. ND6WE1A is a member of a microbial consortium at the cathode of a solar microbial fuel cell originally enriched in seawater, New Jersey, USA. *R. aerolata* CCM 8669 is a type strain from the microbial culture collection in Beijing, China. *R. terrae* KJZ-14 and *R. amarae* KJZ-9 were isolated from soil dirt in Minhang District of Shanghai, China, and *R. nasimurium* E1706032 was isolated from a bird brain at a duck farm in Jining, Shandong Province, China. The adaptation of these strains to life in such contrasting ecological niches is reflected in their species diversity (AAI < 90%) within the genus *Rothia*, to which they were originally assigned by their authors.

It should be noted that according to the results in Figs. 3–7, *Rothia* sp. ND6WE1A, which represents Isolate SG at the whole genome level, and *R. dentocariosa* ATCC17931^T^, which represents the type species of *Rothia* (Austin, 2015), are members of different monophyletic groups separated by intergeneric AAI values < 65%. Thus, when the strains belonging to such groups are appropriately verified (Figs. 3–7), priority in the genus name *Rothia* should be given to the cluster with a member of the type species *R. dentocariosa*. Hence it follows that the strains comprising cluster a in Figs. 3–7 (including *Rothia* sp. ND6WE1A) should be assigned a new genus name that would differ from those in cluster b (Figs. 3–7). This is consistent with the results of the MiGA-based test for *Rothia* sp. ND6WE1A, which showed its statistically valid taxonomic novelty in relation to the TypeMat reference database, containing type whole-genome material. Additional physiological–biochemical and genetic studies are required to resolve this issue with respect to Isolate SG (Sokolov *et al*., 2021).

In the AAI-profiler test (Fig. 8), *Rothia* sp. ND6WE1A, the reference strain for Isolate SG, was the same species as *R. amarae* KJZ-9, with ANI and AAI values of about 99%. That they live in largely different ecological niches (see above) is very probably ensured by the phenotypic traits controlled by the genes in the accessory part of the pangenome. These genes, introduced by HGT (Koonin, 2012), are not accounted for in the tests based on 16S rRNA, ANI, AAI, GTDB-Tk, and MiGA (outside the “MyTaxa Scan” section), which handle “housekeeping” genes. Consequently, if a polyphasic approach is used for these strains (Oren and Garrity, 2014), which takes both genotypic and diverse phenotypic traits into consideration, one can expect that the level of their evolutionary proximity and their taxonomic qualification will be revised substantially (Shchyogolev, 2021).

Our results show that the introduction into consideration in (Parks *et al*., 2020) of the ANI and AF, which ensure quantitative identification of OTUs at the species level for the GTDB bacterial and archaeal genomes (Genome Taxonomy Database, https://gtdb.ecogenomic.org), solves the problem of constructing a taxonomic structure from domain to species (Parks *et al*., 2020) only in part. This is due to the contradictions identified by us in the genus names of strains and metagenomic objects in GTDB R06-RS202 (GTDB Data, https://data.gtdb.ecogenomic.org/releases/release202/202.0/). These contradictions, however, can be eliminated with the AAI-based test (Rodriguez-R and Konstantinidis, 2014; Goris *et al*., 2007; Medlar *et al*., 2018), with appropriate reclassification of the objects.

The application of phylogenetic taxonomic procedures improves the classification of bacteria, but even so, the need remains to further clarify the relationships within the taxon, which encompasses organisms of agricultural, biotechnological, clinical, and ecological importance (Nouioui *et al*., 2018; Xiao *et al*., 2023).

## Footnotes

1 Abbreviations: AAI – average amino acid identity; AF – alignment fraction; ANI – average nucleotide identity; GTDB – genome taxonomy database; HGT – horizontal gene transfer; MF – matched fraction, coverage; MiGA – microbial genomes atlas; MLSA – multilocus sequence analysis; MSA – multiple sequence alignment; OTUs – operational taxonomic units; REQ – rate of elementary quartets.

## ACKNOWLEDGEMENT

We thank Dmitry N. Tychinin (IBPPM RAS) for translating the original manuscript into English.

